# Validation of melomind™ signal quality: a proof of concept resting-state and ERPs study

**DOI:** 10.1101/2020.02.28.969808

**Authors:** Giuseppe Spinelli, Alexandre Odouard, Marie-Cécile Nierat, Sébastien Campion, Mickael Bensoussan, Fanny Grosselin, Katerina Pandremmenou, Audrey Breton, Mathieu Raux, Yohan Attal, Thomas Similowski, Xavier Navarro-Suné

## Abstract

Wearable EEG systems have become accessible to researchers and clinicians over the last decade, thus requiring neurotechnology companies to seek for outstanding EEG signal quality. Here, we show that the melomind™ headset equipped with dry electrodes (myBrain Technologies, Paris, France) allows the recording of reliable electro-cortical dynamics as compared to a wet-based standard-EEG system (actiCAP, Brain Products GmbH, Gilching, Germany). EEGs were acquired simultaneously from the two systems while thirteen subjects underwent a visual oddball, a steady-state visually-evoked potentials (SSVEPs) and two resting-state (RS) tasks. RS were acquired with eyes-closed and eyes-open (2 minutes each) and repeated twice (before and after the cognitive tasks). During the oddball task, participants responded on a gamepad when a target-stimulus was displayed. In the SSVEPs, visual responses were elicited at 15 and 20 Hz through a series of 15-second stimuli presented 5 times each. The power of theta- [4-8 Hz], alpha- [8-13 Hz], and beta- [13-30 Hz] band was extracted from the two RS. The signal-to-noise-ratio in the 15 (± 1) and 20 (± 1) Hz range was computed from the SSVEPs. The shape of the N2/P300 complex was analysed from the oddball task. Strong correlations resulted between the parameters obtained from the two EEG systems (0.53 < Pearson’s r < 0.97). Bland and Altman analysis revealed small dissimilarities between the two systems, with values laying in the 95% confidence interval in all the tasks. Our results demonstrate that the melomind™ is an affordable solution to reliably assess humans’ electro-cortical dynamics at-rest and during cognitive tasks, thus paving the way to its use in neuroscience studies and brain-computer interfaces.

## Introduction

Recent advances in engineering and new discoveries in neurosciences have led to the birth of numerous neurotechnology companies aimed at interfacing the human brain to the machines. As a consequence, a broad number of low-cost, user-friendly devices has appeared on the market over the last decade. Among these, wearable electroencephalography (EEG) systems have increasingly attracted attention from both researchers and clinicians for their application potential (e.g., brain-machine interfaces, gaming, simulation), their extremely flexible utilization (Mihajlovic, Grundlehner, Vullers, & Penders, 2015), as well as their relatively low invasiveness as compared to “standard” and largely-more expensive EEG devices.

EEG traces recorded in laboratory settings used to require outstanding data quality in order to allow analyses in multiple domains, such as the event-related potentials (ERPs) in the time-domain, resting-state (RS) or visually-evoked potentials (SSVEPs) in the frequency-domain, brain oscillatory dynamics and connectivity in the time-frequency domain, and beyond. Newer wearable EEG systems are equipped with effective hardware (e.g., amplifiers specifications, electrode quality) and software (e.g., real-time computations, on-line data quality algorithms, e.g., Grosselin et al., 2019) components to achieve this goal (see Lopez-Gordo et al., 2014 for a systematic review). Therefore, researchers are increasingly prone to use them in clinical trials and to disseminate their findings in scientific journals. In this regard, the majority of the studies using portable EEG devices (e.g., Muse, Emotiv EPOC) assessed the quality of the signal via ERPs analyses (Debener, Minow, Emkes, Gandras, & de Vos, 2012; Duvinage et al., 2013; Badcock et al., 2013; Badcock et al., 2015; Krigolson, Williams, Norton, Hassall, & Colino, 2017). More specifically, these works mainly focused on the manifestation of the N2 and the P300 waveforms, two consecutive components of humans’ ERPs which are convincingly thought to reflect attentional resources allocated to visual stimuli (Luck & Hillyard, 1994; Polich, 2007). However, EEG studies do not limit to ERPs, but often include both SSVEPs and RS tasks. SSVEPs are elicited naturally from visual stimulation at specific flickering frequencies and have been shown to serve to many applications in cognitive (e.g., Vialatte, Maurice, Dauwels, & Cichocki, 2010; Zhang, Jamison, Engel, He, & He, 2011 and clinical neuroscience (e.g., Sokol, 1976; Birca, Carmant, Lortie, Vannasing, & Lassonde, 2008), as well as in brain-computer interfaces (e.g., Giove, 2003; Kluge & Hartmann, 2007; Won, Zhang, Guan, & Lee, 2014). Yet, recording the brain activity at-rest (RS) has revealed crucial information on the functional significance of distributed neural network dynamics, on the hierarchy governing their interactions (e.g., Mantini, Perrucci, Gratta, Romani, & Corbetta, 2007; Brookes et al., 2011) and on the “language” these networks speak (i.e., frequencies) to mutually exchange information on perceptual, motor, cognitive and emotional states. Cross-sectional or longitudinal changes of some of the automated EEG parameters extracted from the resting-brain have increasingly been used to assess or - in some cases - predict the onset of neurological diseases, such as the Alzheimer’s (Stam et al., 2005) or the Parkinson’s (Jackson, Cole, Voytek, & Swann, 2019) disease.

To the best of our knowledge, a number of systematic studies exist on the comparison between dry vs. wet electrodes -based portable EEGs (see Lopez-Gordo et al., 2014 for a review), but the extent to which a given portable EEG system can reliably allow the assessment of humans’ electro-cortical dynamics in a wider range of laboratory protocols as compared to a standard, high-quality EEG is still less explored.

In this work, we addressed all these issues by recording individuals’ brain activity using the melomind™ headset (myBrain Technologies, Paris, France; https://www.melomind.com) and the actiCAP (BrainProducts GmbH, Gilching, Germany; https://www.brainproducts.com/productdetails.php?id=68), simultaneously. Participants underwent a comprehensive protocol including two-resting state recordings, a visual oddball task and a SSVEPs task. Signals recorded from the melomind™ were compared to the signals recorded from the actiCAP in order to assess both the efficacy of the melomind™ to acquire the canonical electro-cortical markers underlying these tasks, and - importantly - the degree of (dis)similarity between the two EEG systems.

## Methods

### Participants

Thirteen individuals (n = 13; 5 males; mean age; 31.4; age range: 20-52) took part in the study following informed consent. Participants had normal or corrected-to-normal visual acuity and reported no symptoms/signs of neurological impairments. The study conformed the 1964 Declaration of Helsinki.

### Procedure, Apparatus, Experimental Setup and Stimuli

Participants sat on a comfortable chair in a well-lightened room while dual EEG was recorded through the melomind™ and the actiCAP. First the actiCAP electrode headset was placed and then the melomind™ (Figure 1).

**Figure 1.**
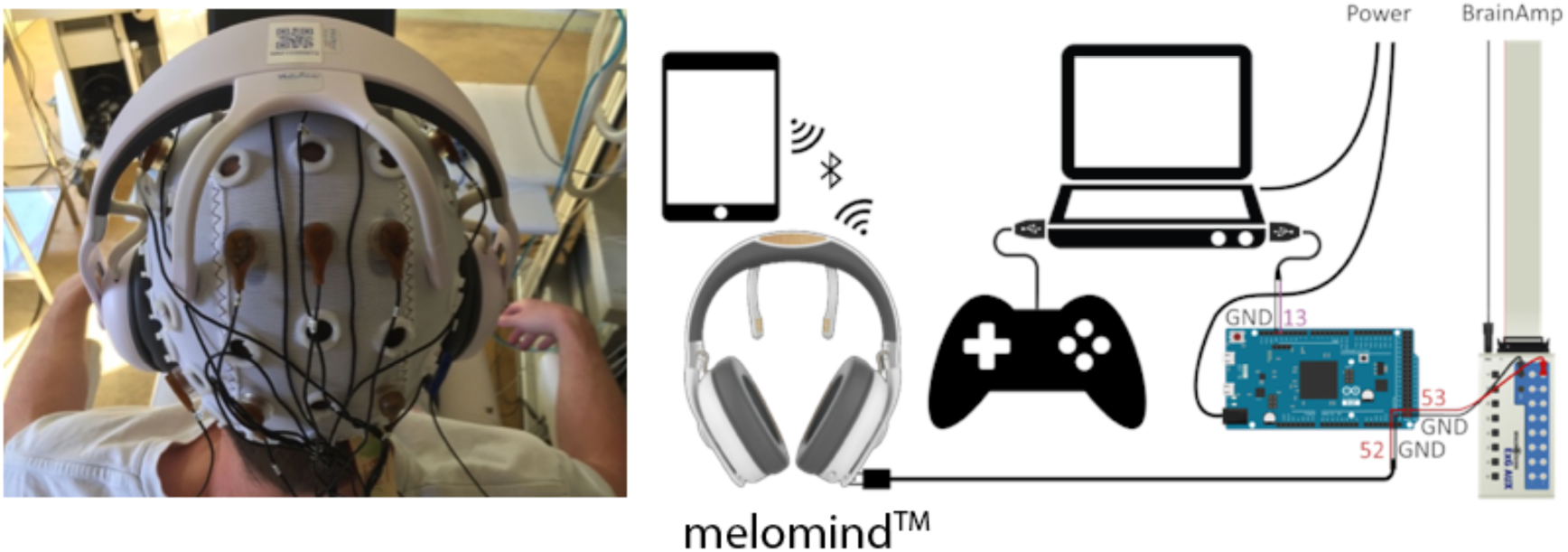
Apparatus and experimental setup. A real recording session with both the melomind™ and the actiCAP (left column). Right-column depicts the experimental setup and the apparatus, including: i) the PC used for stimuli presentation, ii) the gamepad used to record individuals’ reaction-times in the odball task, iii) the Arduino DUE device used to convey and synchronize event-markers to the melomind™ and the actiCAP (i.e., the ExG AUX Box manufactured by Brain Products), iv) the melomind™ headset and v) an example of a portable device to which the signals acquired from the melomind™ where sent via Bluetooth.

The whole experiment lasted about 1 hour and consisted of two Resting State (RS) EEG recordings - one at the very beginning (RS-PRE) and one at the end (RS-POST) of the whole procedure - one visual oddball task (P300 task), and one Steady-State Visually Evoked Potentials (SSVEPs) task. All the tasks, were coded in Python 3.6 using the PsychoPy 3 toolbox (https://www.psychopy.org). Both the source-codes of the tasks and documentations are freely downloadable at https://github.com/mbt-administrator/Research.Tasks.

The order of the oddball and SSVEPs task was balanced across subjects. Stimuli were presented on an ASUS Strix Hero II Laptop equipped with an 15.6-inch LCD screen (refresh-rate: 60Hz) and a GPU NVIDIA® GeForce RTX 2060 (https://www.asus.com/Laptops/ROG-Strix-Hero-II/Tech-Specs). The laptop was placed at a distance of approximately 60 cm from the participant and connected to an Arduino DUE device (https://www.arduino.cc/en/Guide/ArduinoDue) which governed the synchronization of event-markers conveyed to the melomind™ and to the actiCAP. Figure 1 depicts the schema of the experimental setup and the apparatus.

Each RS condition (i.e., RS-PRE and RS-POST) lasted 4 minutes and consisted of 2 minutes eyes-closed and 2 minutes eyes-open EEG recording. Before undergoing the task, participants received written instructions on the PC screen upon the procedure. First, they were asked to close their eyes and keep them closed until a 0.5 Hz alert tone was provided. Then, they were requested to continue with the eyes-open recording by pressing a key on the PC-keyboard. In this phase, a white fixation-cross (against a black-background) was presented on the screen to minimise eyes movements and blinks. Event-markers were sent to the EEGs at the onset, the middle and the offset of each RS recording.

During the P300 task, 10 series of blue (RGB = [0 0 255]) and green (RGB = [0 255 0]) coloured squares (against a black background) were randomly presented for 800 ms. Each stimulus was preceded by a white fixation-cross lasting from 500 to 1500 ms to avoid any habituation effect. The frequency of the blue stimulus (i.e., the ‘rare’ one) differed across the 10 series, such that in 5 series its frequency was 20% (i.e., 6 times over 30 stimuli), and in 5 series its frequency was 40% (i.e., 12 times over 30), with the order of series being completely randomised across subjects. Participants were instructed to press a key-button on a gamepad as soon as the blue stimulus had appeared on the screen. The task counted a total of 300 trials (i.e., 10 series times 30 stimuli) and lasted approximately 20 minutes. A short pause occurred between each trials-series. Event-markers were sent to the EEGs at the onset of each series and at the onset of each stimulus of interest.

In the SSVEPs paradigm, visual responses were elicited at 15 Hz and 20 Hz. This allowed us to eliminate both i) the superposition of the alpha range (8-13 Hz) and ii) any rendering issue due to the 60 Hz refresh-rate of the LCD display. Each subject was instructed to focus on a white square (against a black background) flickering at the stimulation frequency (i.e., 15 or 20 Hz) for 15 seconds. Participants underwent 10 trials, 5 of them at a flickering-stimulation of 15 Hz and 5 at 20 Hz. Each stimulus was preceded by a white fixation-cross (against a black background) lasting 3 seconds and followed by a 45-second pause. A 0.5 Hz tone was delivered 5 seconds before the onset of each trial to alert participants on the onset of the new trial. The order of trials was randomised across subjects. Event-markers were sent to the EEG at the onset, the middle and the offset of each trial.

### EEG recording and analysis

Individuals’ EEG was acquired simultaneously with the melomind™ and the actiCAP (Figure 1). The melomind™ is a portable and wireless EEG headset equipped with two gold-coated dry sensors placed on P3 and P4 sites according to the 10-20 International System. Two textile electrodes are placed on the left- and right-mastoids which serve as ground and reference sites, respectively. EEG signals are acquired at 250 Hz and sent via Bluetooth to a mobile device for both visualisation and storage purposes. As for the actiCAP, signals were acquired from 11 wet electrodes (i.e., F3, Fz, F4, C3, Cz, C4, P1, Pz, P2, O1, O2) arranged on the scalp according to the 10-20 International System. The AFz served as ground electrode and P10 as online reference electrode. The P10 electrode site was chosen as reference electrode because of its closer proximity to the melomind™ reference location (i.e., the right mastoid). The signals were amplified and digitalised at 1000 Hz and then down-sampled offline at 250 Hz to allow comparison with the melomind™. Impedances were kept below 25 KOhm, according to the factory standards for achieving outstanding data quality when using active electrodes. Event-markers were sent simultaneously to both the auxiliary port of the melomind™ (mini-USB) and the Brain Product auxiliary device (i.e., ExG AUX Box, with 8 bipolar inputs and 8 AUX inputs) via a custom-made circuit governed by an Arduino DUE device and controlled by *ad hoc* PsychoPy 3 routines (https://www.psychopy.org/api/serial.html). Baud-rate of the serial port was 9600 Hz. The time taken for each TTL pulse to reach the two systems from the Arduino after any command was sent from the PsycoPy routine was measured by an oscilloscope. In this, a series of TTL (Transistor-Transistor Logic) pulses was sent, resulting in a mean lag of 30 ms. The lag between the two systems to receive a pulse was < 1 ms (range: 0-3 ms), which was mainly accounted for by their different sampling rates.

EEG analyses were performed using both the MNE-Python toolbox (release: 0.18; https://martinos.org/mne/stable/index.html; (Gramfort, 2013); (Gramfort et al., 2014)) and in-house made routines in Python 3.6.

#### Resting-state analysis

For each participant, continuous EEG traces from the melomind™ and from the actiCAP were filtered with a dual pass Butterworth filter of 1-30 Hz and a 50 Hz Notch filter. Eyes-closed and eyes-open segments were then created for the two Resting-State recordings (i.e., RS-PRE and RS-POST), each lasting 2 minutes. A frequency analysis was then carried out by means of a multi-tapered Fast Fourier Transform (FFT) using Hanning taper in the 2-30 Hz frequency-range. Band-power was computed by taking the squared magnitude of the real and imaginary Fourier spectra. The relative power was obtained by dividing the power in the eyes-closed condition to the power of the eyes-open condition. The median activity from the bi-parietal electrodes of the melomind™ (i.e., P3/P4) and of the actiCAP(i.e., P1/P2) was taken. Values of the whole power-spectrum (2-30 Hz) as well as the power-spectra in the theta (4-8 Hz), alpha (8-13 Hz) and beta (13-30 Hz) range were extracted from each participant and from both the RS-PRE and RS-POST condition, separately.

#### P300 analysis

A dual pass Butterworth filter in the 0.5-30 Hz range was applied to each individuals’ EEG to avoid any signal distortion in the ERPs (Tanner, Morgan-Short, & Luck, 2015). A 50 Hz Notch was also applied. The EEG traces were first segmented by taking 500 ms before and 800 ms after the onset of each stimulus of interest and then corrected with respect to the 500 ms time-window preceding the onset of each stimulus. The resulting epochs were first averaged for each condition and then subtracted each other (i.e., ‘rare’ *minus* ‘frequent’). Finally, the median across the bi-parietal electrodes was computed from the melomind™ (i.e., P3 and P4) and from the actiCAP (i.e., P1 and P2). For each participant, the following features were extracted: i) voltage values (in μV) in the whole epoch (from -500 to 800 ms), ii) voltage values (in μV) in the N2 time-window (from 150 to 250 ms) and iii) voltage values (in μV) in the P300 time-window (from 250 to 550 ms).

#### SSVEPs analysis

Individuals’ neural time-series were filtered using a 2-30 Hz dual pass Butterworth filter, and a 50 Hz Notch filter. Data was then segmented from the onset of the stimulus of interest (i.e., 15 Hz and 20 Hz) to 15 seconds after. A multi-tapered FFT was then computed in the 2-30 Hz frequency range and the band-power was calculated form each subject, electrode and trial. The average across trials was then computed, and the median across the bi-parietal electrodes of the two systems (melomind™ [P3/P4]; actiCAP [P1/P2]) was taken. The Signal-to-Noise ratio (SNR) was calculated to quantify the strength of the visually evoked brain response compared to the background power (Tautan, Serdijn, Mihajlovic, Grundlehner, & Penders, 2013). To achieve this in each condition separately, each power spectra was normalised with respect to the sum of the power-spectrum outside the band of interest (i.e., 15 ± 1 Hz and 20 ± 1Hz). Individuals’ whole power-spectrum (2-30 Hz) was extracted from the two conditions, as well as the power-spectra in both the 15 Hz (± 1 Hz) and 20 Hz (± 1 Hz) range.

In order to assess statistically any (dis)similarity between the features extracted from the melomind™ and from the actiCAP, two approaches were used, namely: i) the degree of agreement between variables, using the Bland and Altman method (Bland & Altman, 1986), and ii) a permutation-based Pearson’s correlation analysis, The Bland and Altman method allowed us to assess the degree of agreement between the melomind™ and the actiCAP and, importantly, the likelihood of disagreement between the two measurements. Noteworthy, this analysis posits that if the degree of disagreement in measuring a given phenomenon between two methods is negligible, one can reliably use the two methods interchangeably. In summary, this method consists in examining the data by plotting their difference against their mean, thus allowing to investigate any relationship between the measurement error (i.e., the difference) and the best estimate of the true value (i.e., the mean). Then, the mean difference between the two methods (*d*) and the limits of agreement, i.e., the 95% confidence interval (*d* ± 1.96 SD) around *d*, are calculated. Good agreement exists if most of the differences lie between the limits of agreement. As for the correlation analysis, the Person’s r coefficient was first calculated for each of the (actual) observed features recorded from the two EEG systems. Then, a permutation distribution of the Pearson’s *r* coefficient was computed by shuffling one of the two arrays of values for *n* times - where *n* is the number of permutations (i.e., n = 1000). The probability of getting a true value was ultimately calculated by means of the Monte-Carlo method.

## Results

Bland and Altman analysis for all the tasks and all the features is presented in Table 1. Table 2 details the results of the correlation analysis.

**Table 1.** Bland and Altman analysis of automated parameters derived from the melomind™ and the actiCAP. The table shows the details of the mean difference between the two EEG systems, and the related limits of agreement (lower- and upper- limit) derived calculated from each task (RS, SSVEPs, and oddball). Values refer to the average of the bi-parietal eletrodes-sites (i.e., P3/P4 for melomind™ and P1/P2 for actiCAP).

| Task | Feature | Mean of the difference | Limits of agreement |  |
| --- | --- | --- | --- | --- |
|  |  | (melomind™; actiCAP) | upper | lower |
| RS-PRE | whole spectrum | -0.46 (1.85; 2.31) | -6.48 | 5.55 |
|  | theta [4-8 Hz] | -0.19 (2.16; 2.35) | -7.03 | 6.65 |
|  | alpha [8-13 Hz] | -2.08 (2.73; 4.81) | -12.9 | 8.74 |
|  | beta [13-30 Hz] | -0.17 (1.54; 1.71) | -3.05 | 2.71 |
| RS-POST | whole spectrum | -0.33 (1.76; 2.14) | 11.69 | 10.95 |
|  | theta [4-8 Hz] | -0.29 (1.75; 2.01) | -4.84 | 4.25 |
|  | alpha [8-13 Hz] | -1.25 (3.73; 4.98) | -27.36 | 24.86 |
|  | beta [13-30 Hz] | -0.17 (1.29; 1.46) | -2.42 | 2.08 |
| SSVEPs (15 Hz) | whole spectrum | 0.0 (0.01; 0.01) | -0.06 | 0.07 |
|  | 15 ± 1 Hz | -0.0 (0.01; 0.01) | -0.03 | 0.07 |
| SSVEPs (20 Hz) | whole spectrum | 0.0 (0.01; 0.01) | -0.08 | 0.08 |
|  | 20 ± 1 Hz | -0.0 (0.01; 0.01) | -0.02 | 0.02 |
| Oddball | whole epoch (-500 – 800 ms) | 0.35 (1.57; 1.21) | -11.97 | 12.68 |
|  | N2 range (150-250 ms) | -1.29 (-0.32; 0.97) | -17.91 | 15.31 |
|  | P300 range (250-550 ms) | 0.86 (4.9; 4.05) | -18.78 | 20.5 |

**Table 2.** Correlation analysis between melomind™ and actiCAP signals. Pearson’s correlation analysis computed between the two EEG systems in each task (RS, SSVEPs, and oddball). The third column shows the correlation coefficient (Pearson’s r) and the related p-value (i.e., * = p < 0.05; ** = p < 0.01; *** = p < 0.001).

| Task | Feature | Pearson's r |
| --- | --- | --- |
| RS-PRE | whole spectrum | 0.86 *** |
|  | theta [4-8 Hz] | 0.58 ** |
|  | alpha [8-13 Hz] | 0.85 *** |
|  | beta [13-30 Hz] | 0.53 *** |
| RS-POST | whole spectrum | 0.95 *** |
|  | theta [4-8 Hz] | 0.87 ** |
|  | alpha [8-13 Hz] | 0.93 *** |
|  | beta [13-30 Hz] | 0.82 *** |
| SSVEPs (15 Hz) | whole spectrum | 0.86 *** |
| | 15 ± 1 Hz | 0.97 *** |
| SSVEPs (20 Hz) | whole spectrum | 0.85 *** |
| | 20 ± 1 Hz | 0.96 *** |
| oddball | whole epoch (-500 – 800 ms) | 0.80 *** |
|  | N2 range (150-250 ms) | 0.91 *** |
|  | P300 range (250-550 ms) | 0.56 *** |

### Resting State data

Figure 2 depicts the power-spectral density (PSD) graphs computed from the melomind™ and from the actiCAP in the two Resting-State conditions (RS-PRE and RS-POST). As expected, the normalisation between the eyes-closed and the eyes-open condition resulted in an increase of the alpha-band (8-13 Hz) power.

**Figure 2.**
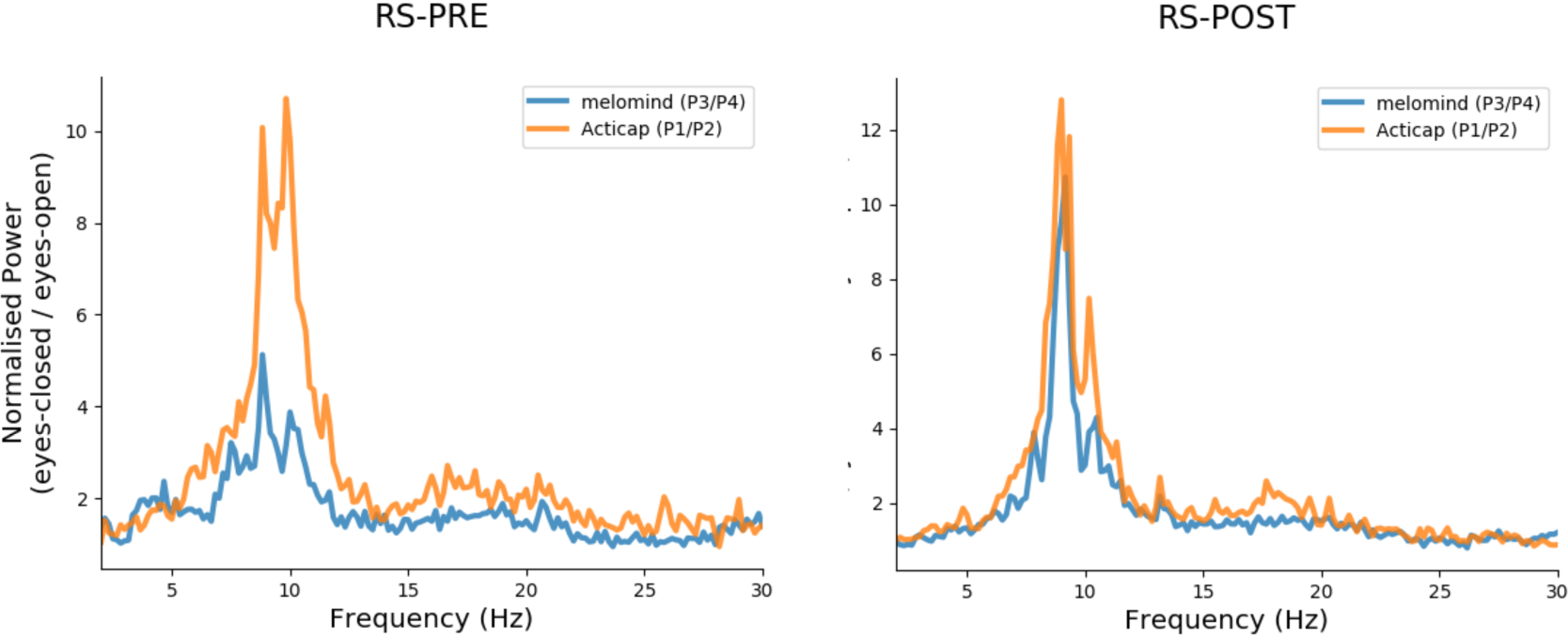
Power Spectral Density graphs of the Resting-State recordings. Figures show the comparison of the PSD between the melomind™ (in blue) and the actiCAP (in orange) in the two resting-state conditions, i.e., RS-PRE (left-column) and RS-POST (right-column). Values refer to the ratio between the eyes-closed and the eyes-open condition. Frequencies (from 2 to 30 Hz in 0.16 linear steps) are plotted on the x-axis.

Bland and Altman analysis for the whole spectrum showed comparable results for the melomind™ and actiCAPin both the RS-PRE (mean difference: -0.46) and RS-POST (mean difference: -0.33) condition (Figure 3). This evidence was replicated for all the canonical frequencies of interest (Table 1) - theta (4-8 Hz), alpha (8-13 Hz), beta (13-30 Hz) - with differences being on average even closer to zero in the RS-POST (overall mean difference: -0.51 ± 0.43) as compared to the RS-PRE (overall mean difference: -0.73 ± 0.79).

**Figure 3.**
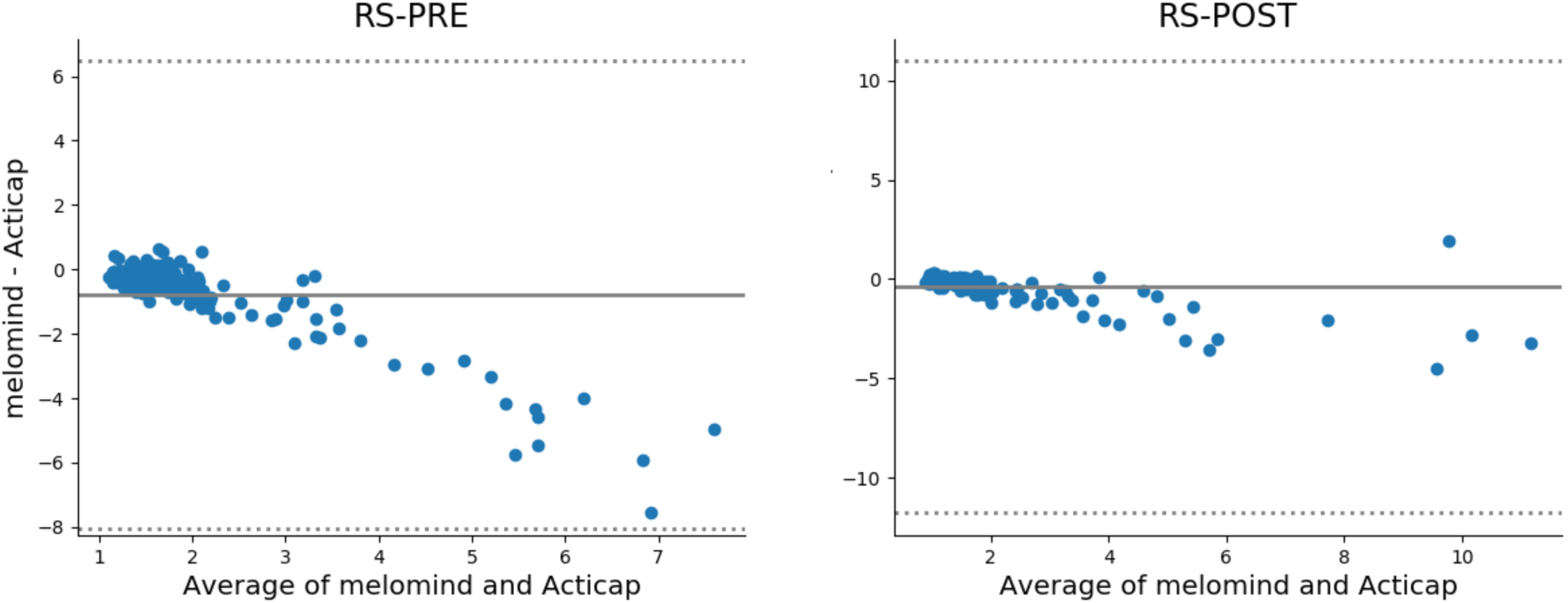
Degree of agreement between the melomind™ and the actiCAP in the RS conditions. The Bland and Altman plot of the bi-parietal electrode-sites (i.e., P1/P2 for actiCAP and P3/P4 for melomind™) computed from the whole power-spectrum in the RS-PRE (left-column) and RS-POST (right-column). Averaged values between the melomind™ and the actiCAP (x-axes) are plotted against their difference (y-axes). Solid-grey lines show the mean difference between the two EEGs. Dotted-grey lines depict the 95% confidence interval around the mean, i.e. the upper- (mean + 1.96 SD) and the lower- (mean - 1.96 SD) limits of agreement.

In addition, the correlation analysis proved a very strong similarity between the melomind™ and the actiCAP, with all the features of interest showing a correlation coefficient higher than 0.5 in the RS-PRE and higher than 0.8 in the RS-POST (Table 2).

### P300 data

Figure 4 depicts the grand mean of the N2/P300 complex as resulted from the melomind™ (in blue) and the actiCAP (in orange). The comparisons between the two signals in the whole epoch-range (from -500 to 800 ms) revealed very small dissimilarities between the melomind™ and the actiCAP, with differences being close to zero (mean difference: 0.35 μV) and lying in the limits of agreement. This result was replicated in the N2 (150-250 ms; mean difference: -1.29 μV) and in the P300 (250-550 ms; mean difference: 0.86 μV) time-window (Table 1). Also, high similarities between the two signals were found, as resulted from the permutation-based correlation analysis. Very-high correlation coefficients were found both in the whole epoch (r = 0.80) and in the N2 range (r = 0.91). A high correlation was rendered in the P300 range (r = 0.56).

**Figure 4.**
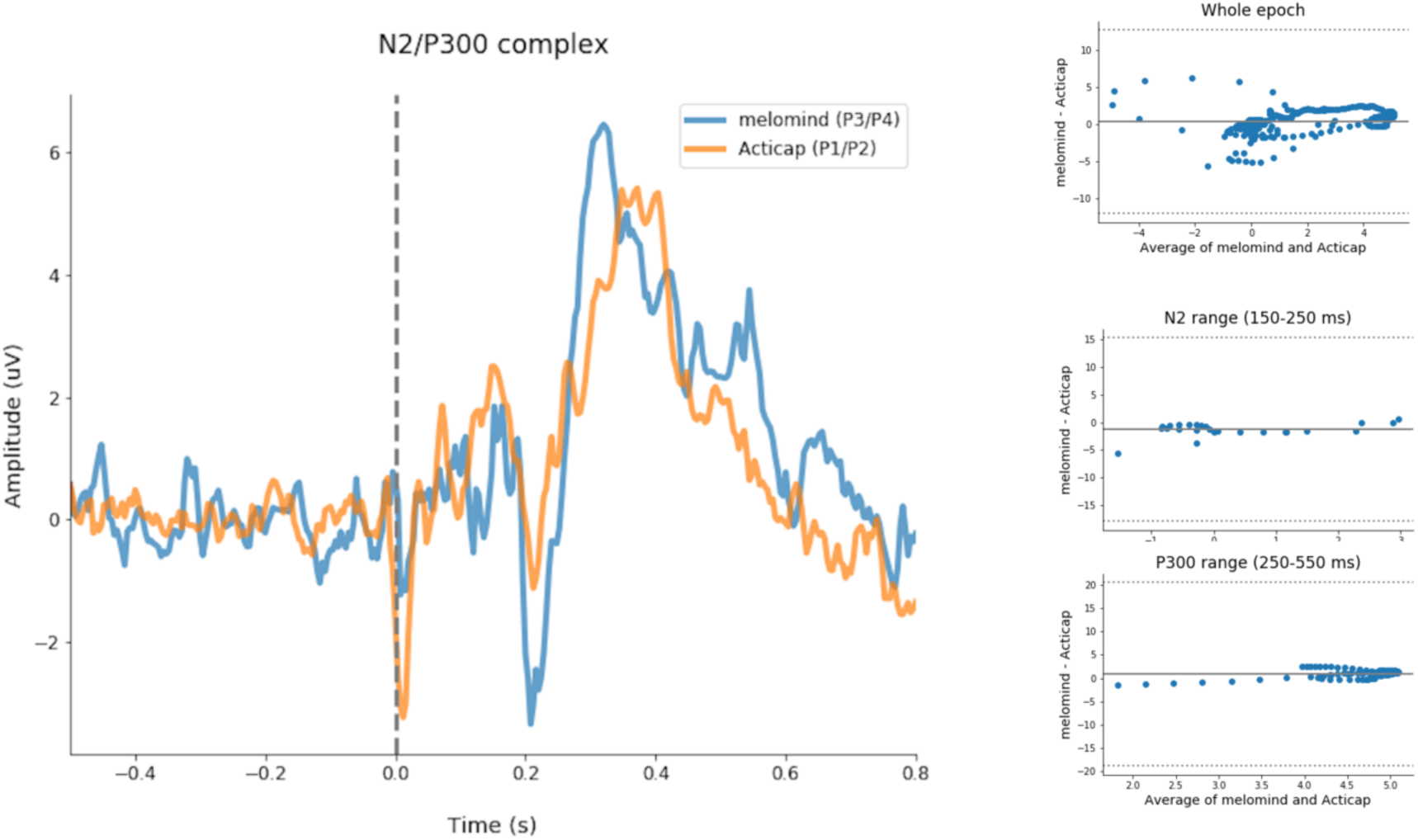
The N2/P300 complex and the agreement between the melomind™ and the actiCAP. Left column depicts the ERPs recorded from the melomind™ (in blue) and from the actiCAP (in orange). The neural time series (in μV) refer to the difference between “rare” and “frequent” stimuli. The signals are corrected with respect to a baseline time-window of 500 ms prior to the onset of each stimulus. The N2 and the P300 components are visible in the time-range from 150-250 ms and 250-550 ms, respectively. Right-column shows the results of the Bland and Altman analysis performed in the whole-epoch range (upper-row), in the N2 range (middle-row), and in the P300 range (lower-row).

### SSVEPs data

The comparison of the power spectral density graphs computed from the melomind™ and the actiCAP in the two-stimulation conditions (15 Hz and 20 Hz) is shown in Figure 5. Noteworthy, the two EEG systems responded adequately to the brain changes induced by the two flickering stimuli.

**Figure 5.**
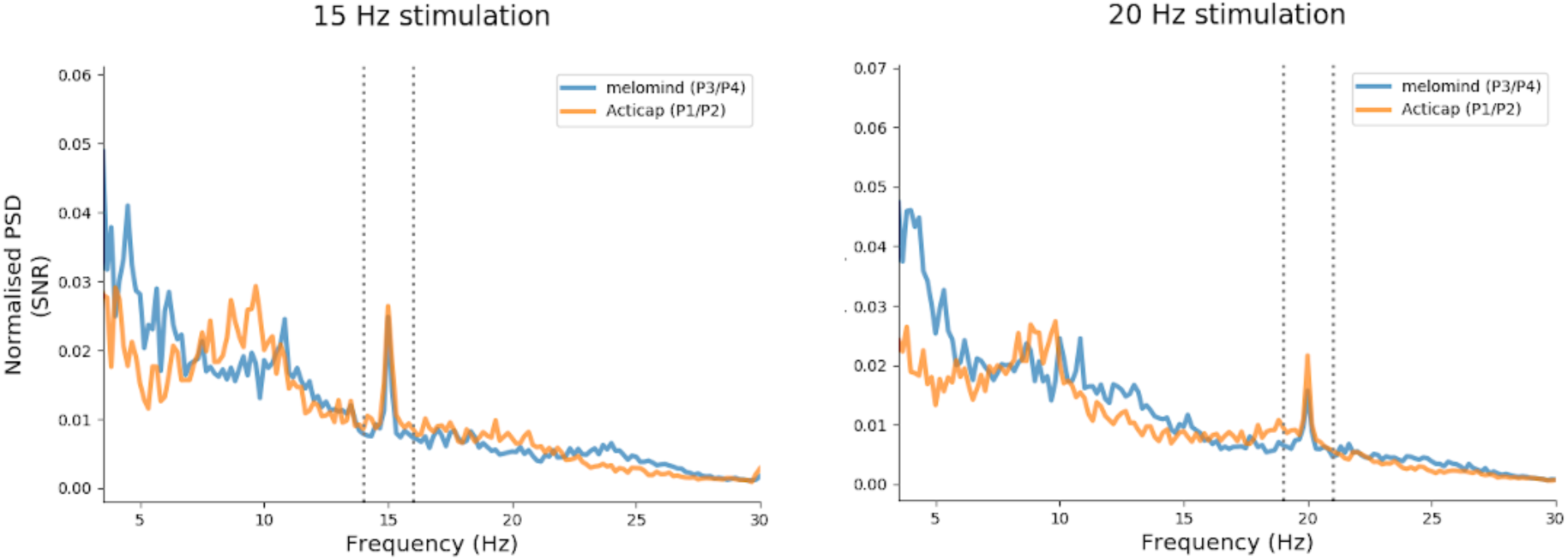
Power Spectral Density maps of the SSVEP task. Figures show the comparison between the melomind™ (in blue) and the actiCAP (in orange) in the two SSVEPs tasks, i.e., 15 Hz (left-column) and 20 Hz (right-column). Power spectra are expressed as SNR. Frequencies (from 3 to 30 Hz) are plotted on the x-axis. The two dotted grey lines depict the frequency-ranges concerned in the analyses, i.e 15 ± 1 Hz and 20 ± 1 Hz.

More specifically, the Bland and Altman analysis showed overall highly comparable power spectra, with differences being extremely close to 0 in both the experimental conditions. These results replicated when considering narrower band-power values around the two frequencies of interests, i.e., when extracting power-spectra in the 14-16 Hz (for the 15 Hz condition) and in 19-21 Hz range (for the 20 Hz condition). In this, the analysis revealed that while the averaged differences between the melomind™ and the actiCAP were more spread around zero (i.e., for the 15 Hz condition: - 0.004; for the 20 Hz condition: -0.003), they still largely lied in the limits of agreements (Figure 6 and Table 1).

**Figure 6.**
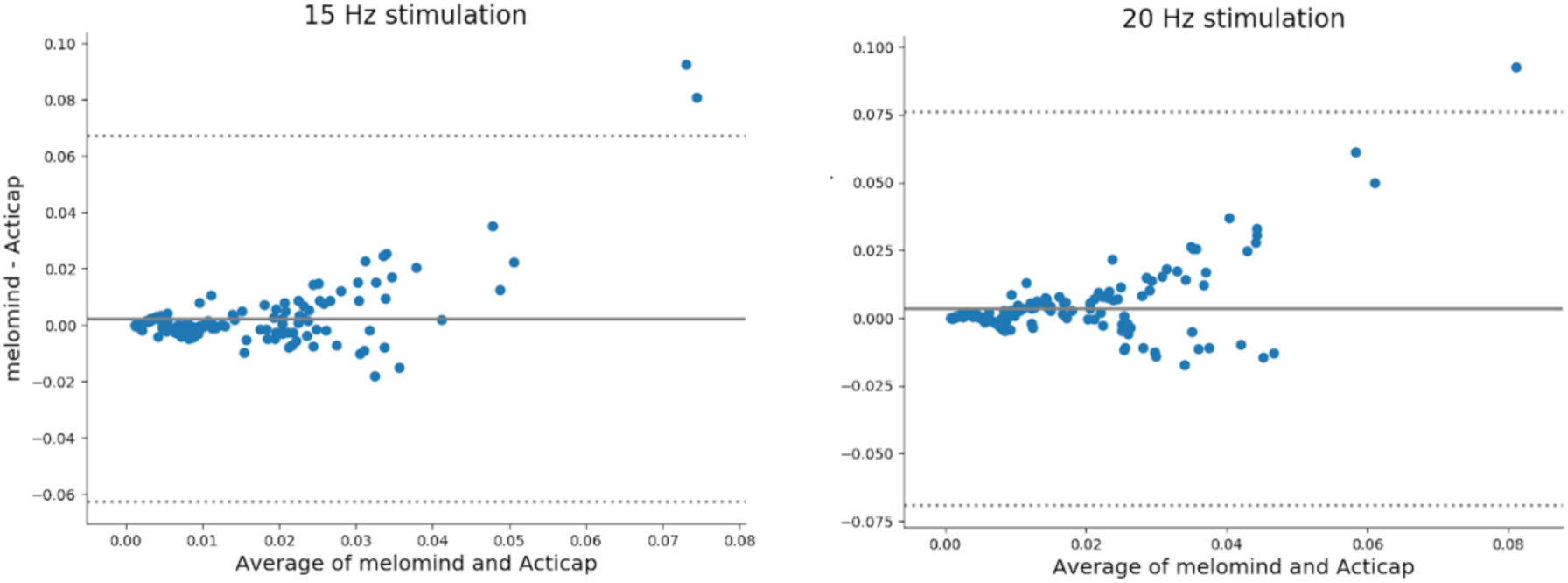
Degree of agreement between melomind™ and actiCAP in the SSVEPs task: The Bland and Altman plot of the bi-parietal electrode-sites (i.e., P1/P2 for actiCAP and P3/P4 for melomind™) computed in both the 15 Hz (left-column) and 20 Hz (right-column) stimulation condition. Averaged values between the melomind™ and the actiCAP (x-axes) are plotted against their difference (y-axes). Solid-grey lines show the mean difference between the two EEG systems. Dotted-grey lines depict the 95% confidence interval around the mean, i.e., the upper-(mean + 1.96 SD) and the lower-(mean – 1.96 SD) limits of agreement.

Moreover, the permutation-based correlation analyses pointed out a very strong similarity between the cortical responses recorded from the melomind™ and from the actiCAP, with all Pearsons’s coefficients being higher than 0.84 (Table 2). Noteworthy, the strength of the correlations increased for the band-frequencies of interest (for 15 Hz: 0.96; for 20 Hz: 0.97) as compared to the whole-spectrum analysis (for 15 Hz: 0.86; for 20 Hz: 0.85).

## Discussion

In this study, we tested the efficacy of the melomind™ headset to record humans’ electro-cortical activity in a laboratory setting. EEG traces were acquired simultaneously from the melomind™ and from the actiCAP while participants performed both resting-state and cognitive tasks. Such a comprehensive approach allowed us to test the behavior of the melomind™ in different experimental protocols, as well as to evaluate any dissimilarities with respect to a standard, high-quality EEG system. In summary, electro-cortical signatures extracted from the melomind™ were comparable and highly correlated to those of the actiCAP in all the tasks.

As for the resting-state recordings, differences in automated spectral parameters were negligible in both the two recordings (i.e., PRE and POST). The RS-POST condition showed closer similarities between the two EEG systems as compared to the RS-PRE. While we do not have a firm examination on this issue, we tend to suggest that dry-electrodes need a wider time to produce data quality as good as a standard-EEG system. In this vein, the “time-factor” might have played a crucial role to ensure a better contact between the dry-sensors and the scalp. Alternatively, one can assume that a wider time span is needed in order for the melomind™ headset to optimally fit (and coexist) with the actiCAP. While on the one hand this is a potential source of limitation of our study, on the other we bring new insights on how simultaneous EEG recording using two different systems (dry-passive vs. wet-active) - which remains likely critical (Lopez-Gordo et al., 2014) - might be implemented to minimize any source of bias. However, the whole power spectrum, the theta- and the beta-band power showed the lowest dissimilarities between the two EEG devices in both RS-PRE and RS-POST. While, on the one hand, differences in the alpha-band power were found to be overall bigger as compared to all the other bands, on the other hand they became sensibly closer to zero in the RS-POST with respect to the RS-PRE. Despite this, similarities between the two systems were optimal, as demonstrated by the permutation-based correlation analysis. This suggests that the melomind™ and the actiCAP are genuinely interchangeable to record brain activity at-rest, albeit i) the analyses were carried out at slightly different bi-parietal sites (P1/P2 vs. P3/P4) and ii) the types of electrodes-set differed between the two devices (passive-dry electrodes for the melomind™ vs. active-wet electrodes for the actiCAP). On a related issue, a previous work showed an overall good correlation (r > 0.48) between signals recorded from dry *vs.* gel electrodes in both eyes-closed and eyes-open resting-state (Tautan, Serdijn, Mihajlovic, Grundlehner, & Penders, 2013). Our results fit with this existing evidence and provide further support on the fact that signal differences can be minimized when similar recording configurations are concerned between two EEG systems, i.e., shorter distances between electrodes. Moreover, here we provide further details on the performance or dry-passive *vs.* wet-active electrodes in recording the power of different frequency-bands. Indeed, communication between two (or more) brain systems is rhythmic (oscillatory) and depends on the “state” of the systems, being a “state” a specific configuration of a system (e.g., hypo/hyper-activation, drowsiness, meditation, relaxation, etc.). Noteworthy, the speed by which brain systems change their states (i.e., the frequencies) are essential to transmit multiple channels of information at the same time across brain networks. In addition, neuroscience studies have shown that spontaneous brain oscillatory activities are of crucial interest to study neuro-cognitive default modes in the “healthy” brain (Laufs et al., 2003; Jap, Lal, Fischer, & Bekiaris, 2009) as well as in a vast variety of neuro-psychiatric diseases, such as Alzheimer (Stam et al., 2005; Kowalski, Gawel, Pfeffer, & Barcikowska, 2001), Parkinson (Klassen et al., 2011), depression (Gotlib, 1998), anxiety (Thibodeau, Jorgensen, & Kim, 2006), autism (Murias, Webb, Greenson, & Dawson, 2007). In this vein, our results are of crucial importance to assess the extent to which the melomind™ headset may allow the recording of oscillatory brain activity in different frequency-bands and possibly pave the way to these research purposes.

In the oddball paradigm, the pattern of the whole evoked potential in the full time-window of interest was comparable between the two EEG systems. No lag was found between the two signals along the time course, thus suggesting that the melomind™ could not only detect fine-grain brain dynamics in the time-domain, but also that the hardware allowed a reliable synchronization of events-markers to the recorded signals. Noteworthy, this is an essential requirement for time-lock analyses, in both the time-domain and the time-frequency domain. Also, in keeping with the seminal validation study of dry and gel electrodes for P300-based BCI (Guger, Krausz, Allison, & Edlinger, 2012), dry-passive electrodes embedded in the melomind™ did not show a different efficacy as compared to a set of wet-active sensors. While previous works performed with wearable EEG systems (Debener, Minow, Emkes, Gandras, & de Vos, 2012; Badcock et al., 2013; Badcock et al., 2015; Duvinage et al., 2013; Krigolson, Williams, Norton, Hassall, & Colino, 2017) showed merely the presence of the N2/P300 complex in the grand-average, here we provided the first direct comparison of the efficacy of a wearable EEG with respect to a standard-EEG in an oddball task. In this, differences between the two EEG systems were negligible and correlations were good for the P300 and high for the N2. P300-based tasks are among the most used ones in neuroscience to investigate humans’ attention. Particularly, the amplitude of the P300 has been used to assess physiological changes of attentional processes (Donchin, Kubovy, Kutas, Johnson, & Tterning, 1973; Polich, 2007), as well as to assess the risk to develop some diseases, such as alcoholism (Polich, Pollock, & Bloom, 1994), schizophrenia (Mathalon, Ford, & Pfefferbaum, 2000), depression (Gangadhar, Ancy, Janakiranaiah, & Umapathy, 1993), or Alzheimer’s disease (Polich, Ladish, & Bloom, 1990). Similarily, the N2 component has been widely used to assess visual attention in healthy subjects (Knight, 1997) as well as for diagnostic purposes in Alzheimer’s (Bennys, Portet, Touchon, & Rondouin, 2007), ADHD (Tye et al., 2014) or binge-eating disorders (Leehr et al., 2018). Also, recent advances in translational neuroscience have led to the birth of P300-based BCI to provide people suffering from motor diseases with a flexible tool to interact quite autonomously with the environment with the hope to recover from their state (Mak et al., 2012; Spüler et al., 2012). While in the present work we did not directly test these issues, our results lead us to suggest that the melomind™ headset may be a suitable device for these purposes.

Finally, the melomind™ and the actiCAP behave almost identically even in the SSVEPs task, with differences being remarkably close to zero in both the two conditions (15 Hz and 20 Hz). Correlations between the two systems showed the overall best results of all the reported analyses, with Pearson’s *r* coefficients being always > 0.79 and the probability of getting a false-positive being always < 0.001. These results demonstrate that the melomind™ can efficiently record sensory brain responses elicited by flickering stimuli at different frequencies, and open the door to SSVEPs-based BCI for both basic (e.g., Wang, Wang, & Jung, 2010; Allison et al., 2012) and translational (e.g., (Ortner, Allison, Korisek, Gaggl, & Pfurtscheller, 2011) research purposes.

## Conclusions

Taken together these results suggest that the melomind™ headset allows the recording of EEG traces as good as a standard, high-quality EEG system (i.e., the actiCAP). Noteworthy, the melomind™ showed an overall outstanding signal quality in a comprehensive experimental protocol, including resting-state recordings and cognitive tasks. The synchronization of marker-events to the continuous EEG signals did not show any lag and allowed the recording of reliable electro-cortical signatures in both the time-(ERPs) and in the frequency (RS, SSVEPs) domain. The high similarities between the electro-cortical signatures recorded from the two EEG systems, together with the high flexibility, the relatively low invasiveness and the low cost of melomind™ pave the way to its use in laboratory settings. Therefore, we encourage the use of melomind™ to further replicate these evidences in a wider variety of cognitive tasks, as well as in different and/or bigger cohorts.

## Acknowledgments

The authors are grateful to all the subjects recruited in the experiment.

## Conflict of interest

myBrain Technologies provides the mobile EEG device used in the present study (melomind™). The authors are employees of myBrain Technologies and had a role in the study conceptualization, methodology, code preparation, data collection, analyses, interpretation and preparation of manuscript

Dr. SIMILOWSKI reports 1) personal fees from ADEP Assistance, AstraZeneca France, Chieisi France, GSK France, Lungpacer Canada, Novartis France, Teva France; and 2) a research grant from Air Liquide Medical System, all unrelated to the present work. Dr. SIMILOWSKI is listed as inventor on issued patents (WO2008006963A3, WO2012004534A1, WO2013164462A1) describing EEG responses to experimental and clinical dyspnea, and his employer has engaged in a contract aiming at exploiting these patients, signed with Air Liquide Medical Systems, France, and myBrain Technologies, France. Dr SIMILOWSKI reports neither fee nor research grant from myBrain Technologies.

## Authors’ contribution

GS, SC, AO, XNS, TS and MR conceived the study. AO and MB implemented the apparatus and the cognitive tasks. GS, MCN and AO acquired the data. GS and AO analyzed the EEG data. GS wrote the manuscript. All the authors discussed the protocol, edited and approved the final manuscript.

